# Detection and Role of Feline Apolipoprotein B mRNA-editing Enzyme Catalytic Polypeptide Subunit 3G-Like Protein in Feline Cells and Tissues

**DOI:** 10.1101/2022.09.03.504613

**Authors:** Barnabe D. Assogba, Shannon Chaudhary, Harmanpreet Kaur, Raymond Soo, Mark A. P. Dela Cruz, Jovy M. G. Assogba

**Author notes:** **Corresponding author:** Dr. Barnabe Dossou Assogba, Ph.D. (principal author), Faculty of Science and Horticulture, Department of Biology, Kwantlen Polytechnic University, 12666 72 Ave. Surrey, British Columbia, V3W 2M8.

## Abstract

The innate host defence system is designed to resist pathogenic microorganism infections. Despite the compelling scientific evidence, our understanding of the full potential of the mechanism is still unclear due to the complex interactions between hosts and invaders. We previously reported latency in cat mucosal infected with low-dose cell-associated feline immunodeficiency virus (10^2^ and 10^3^ infected cells). Here we investigated the expression of Apolipoprotein B mRNA-editing enzyme catalytic subunit 3G (APOBEC3G or A3G) in feline cells and tissues and whether its presence antagonizes the viral pre-integration complex resulting in partial or complete FIV latency. Total RNA and protein lysates were collected from cell lines, blood, and tissue samples. Real-time RT-PCR and western blot assays were used to quantify fA3G-like protein in cats exposed to high versus low-dose cell-associated FIV. We consistently detected fA3G-like protein in mock T-cell lines (E-CD4+, MYA-1, Crandell feline kidney cells) and primary bone marrow-derived macrophages with variable expressions in feline peripheral blood mononuclear cells (PBMC). In addition, the fA3G-like protein was found to interact with FIV group-specific antigen (Gag) protein through immunoprecipitation assays. The protein expression was utterly abrogated following FIV infection. However, *in* lytic FIV infection (in vivo), fA3G-like protein decreased in early post-infection, whereas latently infected cats showed stable expression. These data are the first report of the fA3G-like protein expression in felines and its abrogation in lytic but not in latent FIV-infected individuals. These results might provide new insight into the role of fA3G-like protein in the host defence mechanism against retrovirus infections.

## Introduction

The host invasion by pathogens is tightly regulated by the interplay of cellular factors that restrict the invader from spreading and cause disease progression in infected individuals. Understanding the antiviral properties of these factors could reveal the host and microorganism interactions, improve animal models of infection, and identify targets for antiviral therapies. Two intrinsic factors that control the permissiveness of lentiviruses in primate cells were identified and reported as tripartite motif protein (TRIM5α) (1) and apolipoprotein B mRNA-editing enzyme-catalytic polypeptide-like (APOBEC) 3 proteins (1–3).

TRIM5α has become an essential restriction factor in mammals blocking incoming HIV-1 viral capsid (1,4–6). Thus, TRIM5α from rhesus macaques is intensely active against HIV-1, preventing the use of HIV-1 in primate animal models for HIV/AIDS. Nevertheless, due to the substitutions in the C-terminus, TRIM5α cannot restrict HIV-1 in humans (1,7–9).

APOBEC is an array of genes found on human chromosome 22, which includes APOBEC1 (A1), activation-induced deaminase (AID), APOBEC2 (A2), APOBEC3A through H (A3A-H), and APOBEC4 (A4) (10–12). A3 gene has been found in mice (11) A3F and A3Z3 (13) were reported in a feline with BET as a specific inhibitor (14,15). A1 was the first discovered and said to be a catalytic component with the ability to create premature stop codons. These results in the expression of a truncated form of apolipoprotein B lipid-transport protein in gastrointestinal tissues (16). AID initiates immunoglobulin gene diversification with an unclear mechanism in B cells (17,18). The biological functions of A2, A3, and A4 genes were unknown until it was discovered that human A3G (the most extensively studied member of the A3 family) inhibits HIV-1 replication. A3G achieves this by editing the newly synthesized minus-strand viral cDNA, causing G-to-A mutations on plus-strand DNA (2,19–23). A3B, A3F, and A3DE have been shown to inhibit antiviral activities in several retroviruses, including the HIV-1, simian immunodeficiency virus (SIV), hepatitis B virus (HBV), and some mouse mobile genetic elements (2,20,21,24,25). In HIV-1, this restriction is counteracted by viral infectivity factor (Vif) protein that binds to A3G and recruits a specific E3 ligase complex that mediates its polyubiquitination through 26S proteasome (26–28). In addition, Vif can moderately inhibit the synthesis of new A3G protein (26). The combined effects essentially lead to the complete intracellular depletion of A3G. Thus, no antiviral factor can be incorporated into the budding virion, and those virions remain fully infectious.

In the absence of Vif expression, virion encapsidation has been proposed to be a prerequisite for A3G antivirus activity (2,19). Thus, native A3G protein cannot block viral replication and has been reported to be at high molecular mass (HMM). The HMM form of A3G is associated with Arthrobacter Luteus (ALU) and other small human Y RNAs critical for genomic stability in activated cells (29). The scientific community has widely accepted the current proposed deamination activity of A3G. However, there is no report on the number of A3G molecule expressions in a single cell, nor is there clear evidence of whether all these molecules are eventually recruited to HMM in activated cells or if some are still free in the cytoplasm.

Moreover, the recent finding of nonequivalence between two A3G active sites (N- 65-104 and C- 257-295) supported the hypothesis that deamination may not always be required for A3G antiretroviral activity (30). In addition, there is no evidence of feline A3G and whether it plays a role in viral latency. Therefore, we sought to investigate its expression and possible mechanisms in our previously established animal model of latency (31) and examined the expression of A3G protein in latently infected cat groups versus lytic infected groups. A3G-like protein was wildly expressed in feline cell lines, primary cells, and tissue RNA was detected. In the presence of FIV, the protein expression mimics human A3G expression in HIV-infected cells. However, a decreased fA3G-like protein in lytic FIV-infected individuals seems to mimic the Vif-dependent event in human cases. We detected a stable fA3G-like protein in latently infected cats, which is abolished in lytic-infected individuals. Furthermore, fA3G-like protein interacted with FIV-Gag in latently infected groups. In conclusion, native fA3G-like protein may interact with the viral DNA to inhibit its nuclear transport.

## Results

### Detection of A3G-like protein in feline cell lines, primary macrophages, and PBMC

A3G has been identified in various human tissues, including lungs, spleen, testis, ovaries, and blood leukocytes such as T lymphocytes and macrophages (32,33). Our recent report of an animal model of latency in which exposure to a low-dose cell-associated virus plays a role in establishing latency (31) leads us to hypothesize that an existing cellular innate factor may play a role in the viral latency. Therefore, we decided to explore the expression of A3G and whether its presence is related to a possible mechanism that could potentially contribute to viral latency. To this end, naïve FIV-permissive feline cell lines, macrophages, and PBMC from specific pathogen-free female cats were cultured for seven days. These samples were further stimulated with interleukin-2 (IL-2) and human granulocyte-macrophage colony-stimulating factor (hGM-CSF). Each cell lysate was collected to detect A3G-like protein expression by western blot. Human lymphoid cells (CEM X 174) were a positive control (Fig. 1A). A3G-like protein was detected in feline cell lines (fCD4E, MYA-1, CrfK) (Fig. 1B) and feline bone marrow-derived macrophage cells. The expression of A3G-like protein in feline PBMC varied from individual to individual. As seen in some PBMC, A3G-like protein was strongly expressed while weakly expressed in other samples (Fig. 1C). These data suggest the expression of the fA3G-like protein in feline cells, and the protein can be detected by western blot analysis using a polyclonal anti-human A3G (Immunodiagnostics Inc.). This result is the first study reporting the expression of the fA3G-like protein in feline cells since A3F was delineated to be expressed in feline cells (14). Therefore, we presumed it would be of great interest to investigate whether this protein plays novel functions in the viral latency latent process.

**Figure 1.**
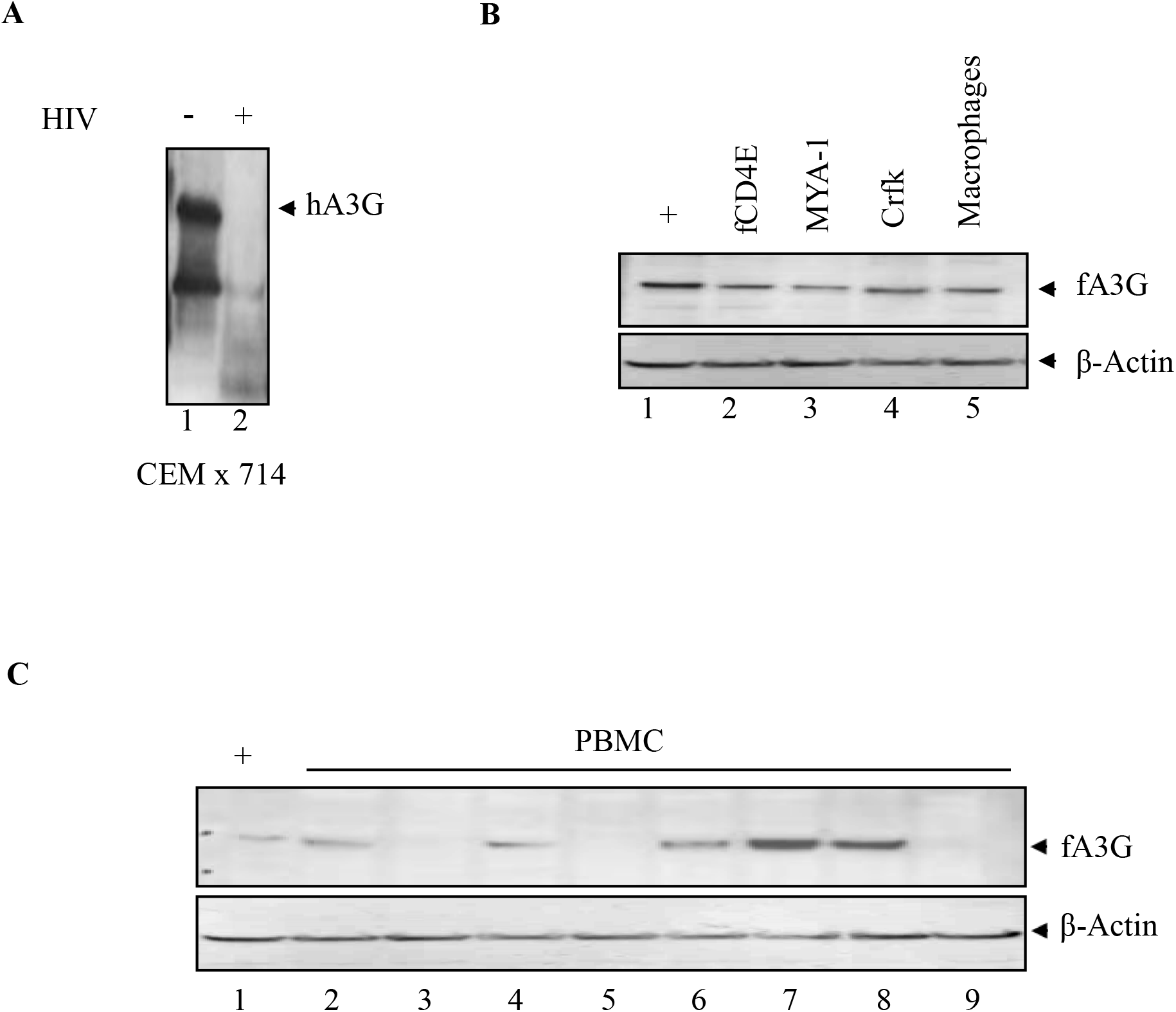
Detection of A3G-like protein in feline cells. (A) Mock and infected human lymphoid cell line (CEM x 714). A3G is expressed in lane 1 but not in lane 2. (B) Naïve feline CD4E T-cells, MyA-1, Crandell kidney cells, and primary feline macrophage Cells in lanes 2, 3, 4, and 5, respectively, lane 1 represents positive control from figure 1 lane 1. (C) Cats PBMC were isolated, purified and total lysates were collected for western blot. The A3G-like protein shows expression in the majority of the primary cells. β-actin was utilized as the loading control.

### Inhibition of A3G-like protein in FIV-infected cell lines and macrophage cells

In blood T lymphocytes, A3G can restrict HIV-1 infection in the virus infectivity factor (Vif)-deficient strains (2). HIV-Vif targets A3G by recruiting elongin (E3) and other cellular factors for ubiquitination and proteasomal degradation (34,35). To investigate whether fA3G-like protein decreased in the presence of FIV-Vif, macrophages, MYA-1, and E-cells were infected with FIV for 72 hours, and total lysates were collected for Western blot assay. fA3G-like protein was detected 24 hours post-infection in E-cells with a decrease at 72 hours post-infection (Fig. 2A). The protein expression was abolished entirely at 72 hours post-infection in MYA-1 cells (Fig. 2B). The protein was wholly abrogated at 48 through 72 hours post-infection (Fig. 2C). The RNA level was determined in both E-cells, MYA-1, and macrophages. fA3G-like-RNA was strongly expressed in mock with a decrease in E-cells at 72 hours post-infection (Fig. 2D). However, the RNA levels were wholly abrogated by the presence of FIV in both MYA-1 and macrophages (Fig. 2E, and 2F).

**Figure 2.**
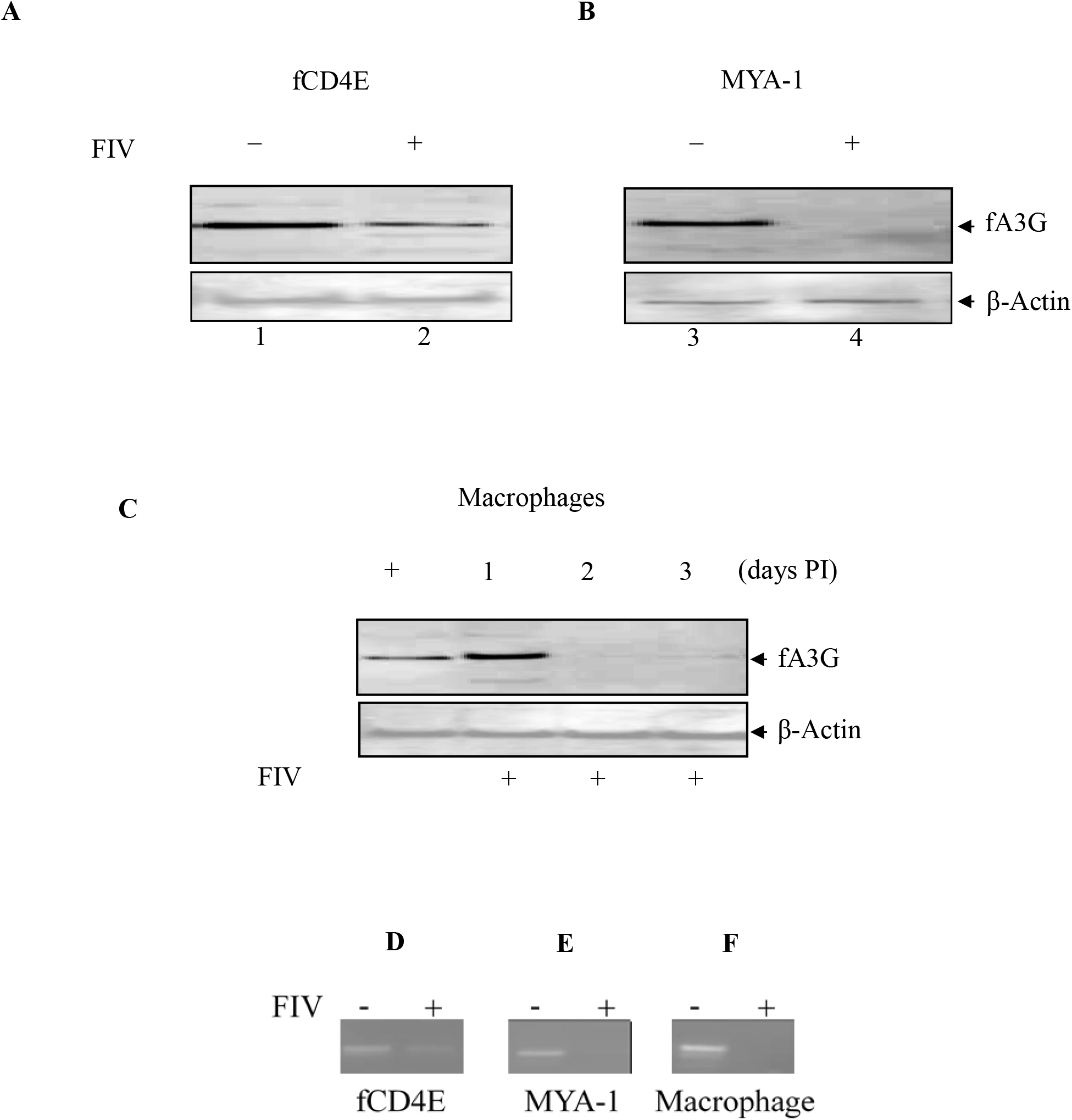
Detection of APOBEC-3G-like protein in FIV infected feline primary and cell lines. The order of samples in (A): lane 1; FCD4E (naïve E-cells), lane 2; infected FCD4E; (B) lane 3, naïve MYA-1; lane 4, infected MYA-1. (C) Lane 1, mock feline macrophages; lane 2, infected feline macrophage at one day post-infection; lanes 3 and 4 represent 2- and 3-days post-infection. fA3G-like gene RNA levels were detected in the above mock cells (lanes 1, 3, and 5) but not in the presence of FIV (lanes 2, 4, and 6) (D, E and F). Data from the β-Actin shows the loading control in the western blot assays.

Vif is a common gene in all lentiviruses, except for the equine infectious anemia virus (36) and has been reported to antagonize A3G in human and simian immunodeficiency viruses through ubiquitination. Although only 32 percent of the HIV-1 sequence is conserved in the FIV gene (37,38), FIV-Vif seems to play a similar role as HIV-Vif in humans. Therefore, we suspected the activity of FIV-Vif and explored whether FIV-Vif plays a role in reducing or eliminating fA3G-like protein in MYA-1-infected cells. Accordingly, we transfected FIV-pF34, pF34ΔBH, or pF34Δ3.4 plasmids into MYA-1 cells for 5 days, and western blots were conducted to detect fA3G-like protein. fA3 G-like protein decreases considerably in FIV-pF34-transfected cells but is not entirely abolished at day 5 post-infection. Interestingly, pF34ΔBH (deletion of 223 pb between *BamHI* position 5482 and *Hind III*, position 5705) and pF34Δ3.4 (with four site mutations) were not able to reduce nor abolish the protein expression in MYA-1 cells (Figure 3A).

**Figure 3.**
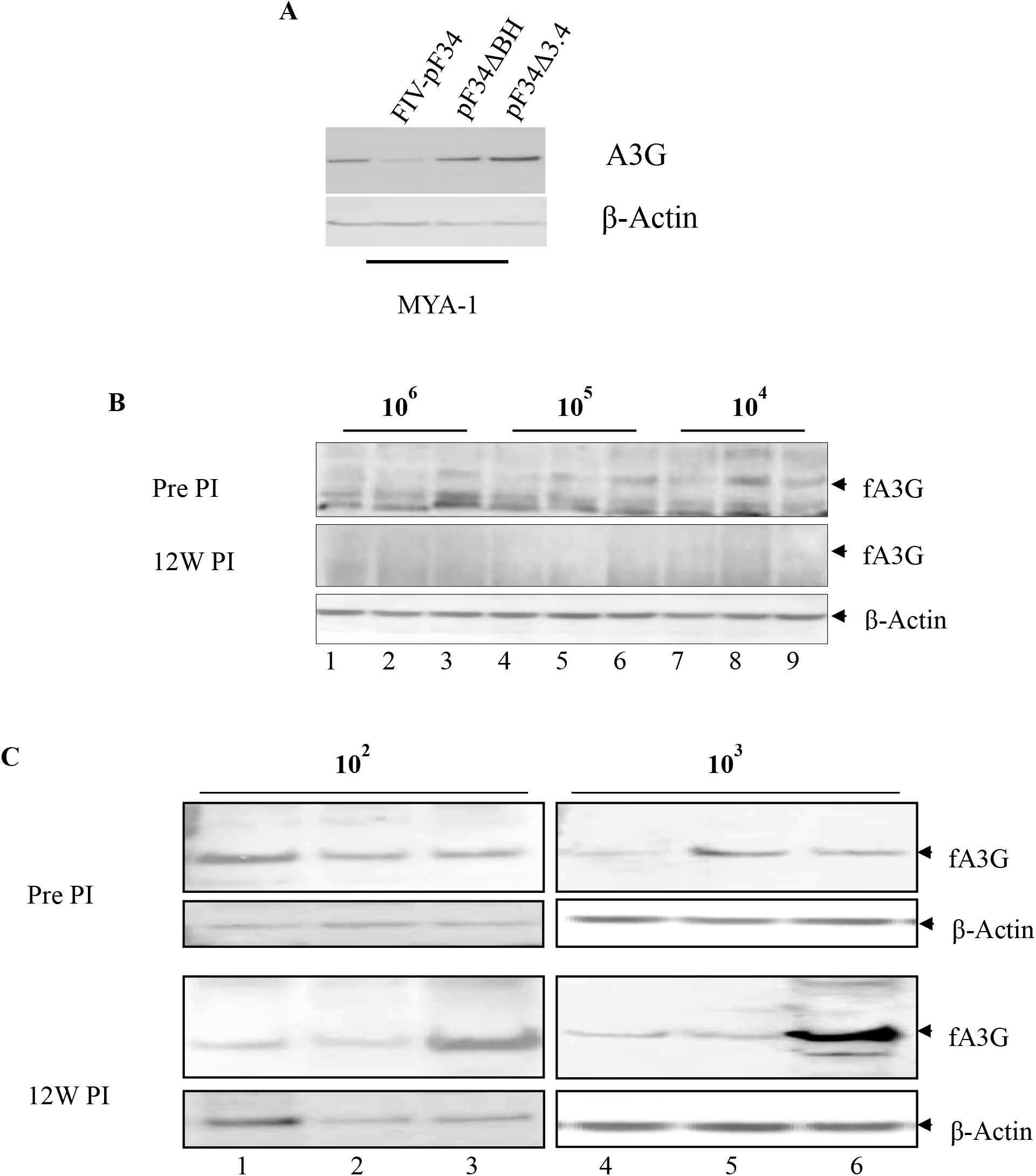
*In vivo* detection of fA3G-like protein in low and high doses cell associated FIV infected feline primary and cell lines and feline Vif monitoring the protein expression. (A) MYA-1 cells transfected with plasmids containing full-size feline Vif in lane 2 (FIV-pF34), lane 3 (pF34ΔBH) and lane 4 (pF34Δ3.4). fA3G-like protein can be seen in lanes 3 and 4 with a faint line in the presence of FIV-pF34 (lane 2). Lane 1 shows positive control. (B) PBMC of lytic infected groups were pre and post-infection tested for expression of the fA3G-like protein. Three cats represented each group, and individuals (cats) were exposed to 10^6^, 10^5^, and 10^4^ FIV-infected cells. The upper film shows the expression of the fA3G-like protein in the PBMC of the group members at the pre-infection stage. The middle film shows the results of the same groups as mentioned above, but these individuals were exposed to FIV, and samples were collected 12 weeks post-infection. The bottom film represents β-actin as the loading control. (C) The same experiment described above was reproduced in individuals exposed to 10^2^ and 10^3^ FIV-associated cells. The last two groups developed latent reservoirs.

### *In vivo* inhibition and stable expression of a fA3G-like protein in FIV lytic versus latent infection

Infection of the transcriptionally active HIV-1 target cells allows a successful reverse transcription and may result in the integration of the viral DNA into the host genome. However, cellular and viral transcription ceases before viral or immunologic cytopathic effects, and the virus becomes dormant. In this form, HIV-1 is resistant to the effects of antiviral drugs and is maintained until the cell is activated (stimulated to increase transcription. The latent stage of HIV-1 has become a significant health issue, posing a formidable obstacle to the goal of eradicating HIV-1 in infected patients. To address whether fA3G-like protein influences FIV latency, we investigated the expression of the fA3G-like protein in both FIV-lytic versus latently infected groups. fA3G-like protein was detected in all groups at pre-infection (Fig. 3B and 3C). However, the A3G-like protein was utterly abolished in lytic infected cats while remaining stable in latently infected groups at 12 weeks post-infection (Fig. 3B and 3C).

### Distribution of fA3G mRNA in a variety of feline tissues and 3’-5’ RACE

To further gain insight into fA3G-like protein expression, we investigated the mRNA level in multiple feline tissues. The fA3G-like protein was broadly expressed in splenocytes, popliteal lymph nodes, iliac lymph nodes, and intraepithelial lymphocytes (Fig. 4). fA3G-like protein was absent in mesenteric lymph nodes and lamina propria lymphocytes (Fig. 4, lanes 4 and 7).

**Figure 4.**
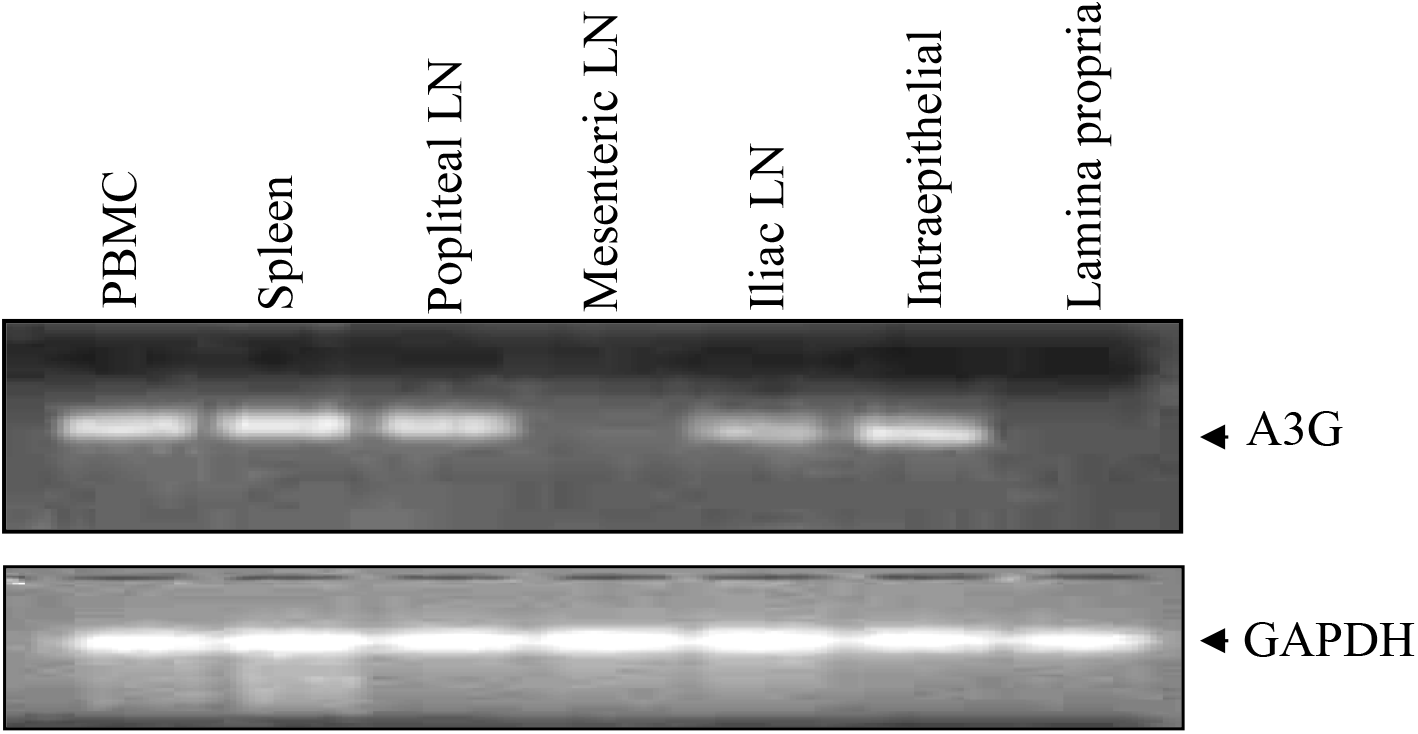
Expression of fA3G-like RNA in multiple feline tissues. Lane 1: PBMC, lane 2: spleen; lane 3: popliteal lymph node; lane 4: mesenteric lymph node; lane 5: iliac lymph node; lane 6: intraepithelial and lane 7: lamina propria. The bottom film shows the glyceraldehyde-3-phosphate dehydrogenase (GAPDH) as the loading control.

To identify the 5′ termini of fA3G transcripts, mRNA was prepared from the fCD4 cell line, followed by calf intestinal phosphatase (CIP) treatment to remove the 5′ phosphates from truncated and non-mRNA lacking a 5′ cap structure. The resulting mRNA was further treated with tobacco acid phosphatase (TAP) to remove the 5′ caps structure. Thus, leaving at the 5’ end, the essential phosphate needed for ligation to the gene racer RNA oligo. An RNA oligo that provides a known priming site for generator PCR primers after the mRNA is transcribed into cDNA was ligated to the 5′ ends of the decapped mRNA. This method of random amplification of cDNA ends (RACE) is highly selective since the CIP treatment of noncapped messages prevents ligating the RNA adaptor to these transcripts. Thus, ligation of the RNA oligo is targeted only to the 5′ ends of full-length transcripts. Using an oligo(dT) primer, the cDNA was generated from the fA3G 5′ adapted mRNA. Reverse transcription (RT)-PCR was then performed using primers specific for the 5′ adaptor oligonucleotide and a reverse fA3G specific primer. The 3′ ends were synthesized by a directly reverse transcript of the mRNA using an oligo (dT) primer and amplified using generator 3’primer conjugated to a forward fA3G specific primer. PCR was conducted in 50 μl using Platinum Taq DNA Polymerase High Fidelity, 25 mM MgCl2, 5 mM dNTPs, 0.5 μM primer, and 35 cycles. RT-PCR products were purified agarose gel and subsequently characterized by DNA sequencing.

### *In vivo* cytoplasmic interaction of the fA3G-like protein with FIV-Gag in latent infection

A successful viral infection is regulated by serial steps rigorously controlled by intrinsic cellular factors designed to block the progression of the disease. These steps included attachment, the release of viral cores into the cytoplasm, core disassembly, the reverse transcription complex, and the establishment of the pre-integration complex (PIC) that will be transported into the nucleus for integration. We suspected the block of the PIC in the cytoplasm by fA3G-like protein. Therefore, we determined whether fA3G-like protein could interact with the pre-integration complex in the cytoplasm of latently infected cells. To detect such interaction, we immunoprecipitated FIV-Gag, and used the same blot to see fA3G-like protein. As a result, FIV-Gag and fA3G-like proteins were expressed with the predicted size (data not shown). Thus, we conclude that a direct interaction between fA3 G-like protein and FIV-Gag protein in the cytoplasm results in the inhibition of PIC nuclear transport.

## Materials and Methods

### Origin of cells and *in vitro* assay

Feline PBMC and tissue lymphocyte cells were collected at necropsy and were maintained in culture as described (31). In addition, FCD4E (feline T-cell line) and feline T lymphoblastoid (MYA-1) Cells were propagated in complete RPMI 1640 media containing 20% fetal bovine serum (Atlanta Biologicals), 1mM sodium pyruvate, 0.1 mM HEPES buffer sodium, 5 × 10^−5^ M β-2-mercaptoethanol, 100 U of penicillin/ml, 100 µg of streptomycin/ml (all from Invitrogen Life Sciences), and 100 U/ml recombinant human interleukin-2 (IL-2) kindly provided by the NIH AIDS Research and Reagent Program at 37°C 5% CO2. In addition, human lymphoid CEM X 174 cells were maintained in RPMI 1640 supplemented with 10% fetal calf serum, two (2) mM glutamine, 0.1 mg of streptomycin per ml, and 100 U of penicillin per ml (Cellgro by Mediatech).

To isolate bone marrow-derived mononuclear cells (BMMC), femurs were removed from a clinically healthy, euthanized, and specific-pathogen-free cat. The femurs were cracked, and the bone marrow was aspirated using 1 x phosphate-buffered saline (PBS). The BMMCs were washed twice with 1 x PBS and cultured in Dulbecco’s modified Eagle’s medium (1.5 × 10^6^ cells/ml) containing 10% FBS, 100 U/ml penicillin, 100 μg/ml streptomycin, two (2) mM glutamine, 50 μM 2-mercaptoethanol for four days, and supplemented once with hrGM-CSF (1000 U/ml) on the first twenty-four (24) hours of culture. After four days, nonadherent cells were removed by two washes with a pre-warmed medium. At seven days, the majority of the adherent BMMCs had macrophage morphology.

### Western blot analysis

Cells were washed three times and lysed on ice with 1 x cell lysis buffer (Cell Signaling) containing 20 mM Tris-HCl (pH 7.5), 150 mM NaCl, 1 mM Na2EDTA, 1 mM EGTA, 1% Triton, 2.5 mM sodium pyrophosphate, 1 mM b-glycerophosphate, 1 mM Na3VO4, 1 μg/ml leupeptin, and 1mM PMSF were added immediately before use as recommended by manufacture (Cell signaling). Twenty-five micrograms of total lysate were mixed with sample buffer (62.5 mM Tris-HCl, pH 6.8, 25% glycerol, 2% SDS, 1% bromophenol blue, and 5% 2-mercaptoethanol), and heated at 90°C for 5 min. The heated samples were briefly centrifuged at high speed before being resolved on 4-12% NuPAGE gel (Invitrogen). Proteins were transferred onto a PVDF membrane filter paper sandwich (Invitrogen) and blocked for 45 minutes at room temperature in a 5% milk solution containing Tris-buffered saline (25 mM Tris, pH 7.6, and 150 mM NaCl). Membranes were overnight probed in 1/1000 Rabbit Anti-Human APOBEC3G (Immunodiagnostics Inc.), washed three times, incubated with HRP-conjugated donkey anti-mouse IgG (1:10,000; Promega), and revealed with Super Signal chemiluminescent substrate reagents (Pierce) by autoradiography.

### DNA and RNA isolation and RACE 5’ ends

PBMC and tissue lymphocyte DNA was isolated as previously described (31). mRNA was isolated using a Micro-FastTrack mRNA kit (Invitrogen). To isolate the full-length fA3G cDNA, 5′-RACE and 3′-RACE were done using the GeneRacer method according to the manufacturer’s instructions (Invitrogen). Briefly, to eliminate truncated and non-mRNA from further analysis, mRNA samples were treated with calf intestinal phosphatase to remove 5’-phosphates. The 5’-cap structure was removed from intact and full-length mRNA by tobacco acid pyrophosphatase, leaving a 5’-phosphate at the 5’-end. The GeneRacer RNA oligo was ligated to its 5’-terminus with T4 RNA ligase. fA3G-specific antisense primer 1 was used to initiate reverse transcription of the 5’-portion of A3G mRNA, including the RNA oligo, with Superscript III reverse transcriptase, which resulted in first-strand cDNA with known 5’- and 3’-priming sites. The 5’-portion of A3G mRNA was amplified by polymerase chain reaction (PCR) with the GeneRacer 5’-primer and fA3G-specific antisense primer 2. The PCR products were gel purified and sequenced. PCR reaction: 94 °C for 2 min, 94 °C for 30 s, 72 °C for 1 min, for 5 cycles; 94 °C for 30 s, 70 °C for 1min, for 5 cycles; 94 °C for 30 s, 68 °C for 30 s, 72 °C for 1 min, for 25 cycles; and 72 °C 10 min for elongation. The PCR products were analyzed on a 0.8% agarose gel.

### RT-PCR and sequencing

All PCR products were excised from 0.8% agarose gel and purified using a QIAquick gel extraction kit (Qiagen) following the manufacturer’s instructions. The purified products were sequenced directly with 3.2 pmol of the appropriate primers. All sequences were assembled using the Seaman program of the Lasergene package, version 6 (DNAStar). Consensus sequences were derived from at least two independent forward and reverse sequences. However, for most of the genome sequence, multiple sequence overlap was achieved using independent PCR products.

Editseq (DNAStar) was used to translate gene sequences and MEGALIGN (DNAStar) to align sequences. The alignments were edited in Genedoc. Percentage identities and similarity scores were determined in the MEGALIGN program. The G+C content was obtained from the Editseq program.

### Plasmids and DNA transfection

The reporter gene plasmids, FIV-pF34, pF34ΔBH, and pF34Δ3.4, were previously described (39) and were generous gifts from Dr. Ellen Sparger (UC Davis School of Veterinary Medicine). Plasmids were amplified in *Escherichia coli* (XL-Blue) extracted by alkaline lysis and purified by column separation (Qiagen). The integrity was confirmed by gel electrophoresis in 0.8% agarose. DNA concentrations were determined by absorbance at 260 nm. According to the manufacturer’s instructions, DNA transfection into fCD4E cells was done using SuperFect transfection reagent (Qiagen).

### Co-Immunoprecipitation

Cell supernatant was used for FIV-gag immunoprecipitation. Protein concentrations were measured using Bradford’s method, and 2 mg of total proteins were used for immunoprecipitation. FIV-gag protein was immunoprecipitated, and interacting proteins were co-immunoprecipitated with 5 µg of anti-gag antibody per mg of total proteins using Dynabeads protein G (Invitrogen). Proteins were solubilized in Laemmli buffer and separated by SDS–PAGE. After electroblotting, nitrocellulose membranes were blocked by TBS, 0.1% Tween 5% milk for A3G Western blotting, and probed overnight at 4°C with the mouse monoclonal anti-IKK antibody (1/2000) or the mouse monoclonal Akt antibody (1/1000), or the rabbit anti-phospho-(Ser/Thr) Akt substrate antibody (1/2000) (Cell Signaling Technology, France). Then, the membranes were rinsed and incubated with peroxidase-conjugated anti-mouse or anti-rabbit immunoglobulin G for 1hr at room temperature. After extensive washes, the reaction was revealed using the Super Signal West Pico chemiluminescent substrate (Pierce, France) with Kodak X-Omat AR film.

## Discussion

This study sought to investigate the expression of A3G in feline cells and tissues and whether its presence antagonizes viral DNA (PIC) for nuclear transport. The data herein reveal the presence of fA3G-like protein in both feline cell lines (Fig. 1B), primary cells (Fig. 1B; lane#5 and Fig. 1C), and tissues (Fig. 4). The protein encloses PIC in the cytoplasm and propels the viral DNA to establish transient or permanent latent reservoirs. FIV-Vif decreases or completely abolishes fA3G-like protein in lytic-infected individuals. We could not explore the mechanism of FIV-Vif quelled fA3G-like protein in feline cells and tissues. However, one prospect of how Vif determines the fate of fA3G-like protein might be through the (40) ability of Vif to direct proteasomal degradation of the fA3G-like protein (40), (41) via ubiquitin E3 ligase complex as reported in humans (42)(43)(44). This study is the first report of fA3G-like protein since fA3F and A3Z3 (13) were found in felines; consequently, our data conflict with previous reports of non-existing fA3G. These results concoct new opportunities for more molecular aspects of research to determine the gene location within feline chromosomes and scour additional physiological functions that this protein might be playing. Detecting the protein in feline cells and tissues was significant attainment that proved once again the importance of small animal models in the fight against HIV. *In vivo* analysis of the protein in cats exposed to a low dose of cell-associated FIV or latently infected groups) indicates the ability of the fA3G-like protein to suppress PIC nuclear localization through direct interaction with FIV-Gag (data not shown).

Several studies have shown the ability of the PIC to gain the nuclear environments by matrix (MA)-mediated nuclear transport, integrase and importinα/β (42), (43). In addition, two other pathways controlled by KIF5B and NUP358 are additional venues for PIC to enter the cell nucleus during HIV infection (45). Herein we are reporting the expression of the fA3G-like protein in feline cells and tissues. In addition, native fA3G can eventually bind to the viral DNA, causing latency in the cell cytoplasm.

The PBMC are blood cells, including lymphocytes, macrophages, and monocytes. These cells are known for their significant contributions to animal defence mechanisms. For example, the B-lymphocytes produce antibodies to mark invaders, antigens, and infected body cells killed by T-lymphocytes. On the other hand, T-lymphocytes defend the body from pathogens by attacking and destroying them. Macrophages are effector cells of the innate immune system that can phagocytose invaders, process, present antigen at their surfaces and initiate immune system activation by producing interleukin-1 (46), (47).

The presence of fA3G-like protein in these cells (Fig. 1C) could imply its association with the immune system to defend the animal body. However, lanes 3, 5 and 9 did not show the expression of the fA3G-like protein. Two different A3G protein forms have been reported; the high molecular mass of 700 KDa (HMM) and Low molecular mass (LMM) ranging from 44-46 KDa were previously reported (48). Thus, we thought that in these individuals, most fA3G-like proteins might be at HMM. Although fA3G-like protein can be seen in mock feline cells and tissues, the protein was either reduced or completely abolished in the presence of FIV (Fig. 2A, 2B and 2C). Vif was previously reported to protect HIV during infection by antagonizing the A3G function. This could justify that we are seeing a reduction or complete eradication of fA3G-like protein and the level of RNA (Fig. 2D, 2E, and D3) in the presence of FIV. Figure 3A supports our hypothesis that feline Vif is the factor that causes fA3G-like protein degradation. The data in figure 2 also strongly supports the *in vivo* pre-infection and 12 weeks post-infection of latently and lytic infection models, respectively (Fig. 3A and 3C).

The decision of FIV to establish latent reservoirs in multiple tissues such as PBMC, spleen, popliteal lymph node, mesenteric lymph node, iliac lymph node and gut tissues through intraepithelial and lamina propria is highly dependent on the dose of the virus that the individuals were suggested to be exposed to (49). fA3G-like protein can be detected in all the above tissues except the mesenteric and lamina propria (Fig. 4). Once again, we might be experiencing the expression of the fA3G at HMM in lymph node and lamina propria, respectively. However, this condition makes detecting fA3G-like protein impossible in our assays.

The nuclear translocation of the viral DNA (PIC) is a complex mechanism that still needs clarification to perceive the true nature of how the viral DNA ends up in the nucleus. The matrix, integrase, importinα/β, Vpr, KIF5B, and NUP358 were all considered critical factors for the DNA’s destination (42), (43), (45), (50). However, by interacting with the viral DNA in the cytoplasm, fA3G-like protein blocks the above factors from having access. Consequently, it stops the nuclear transport of PIC, pressuring the virus to establish latent reservoirs in the cytoplasm (Fig. 5).

**Figure 5.**
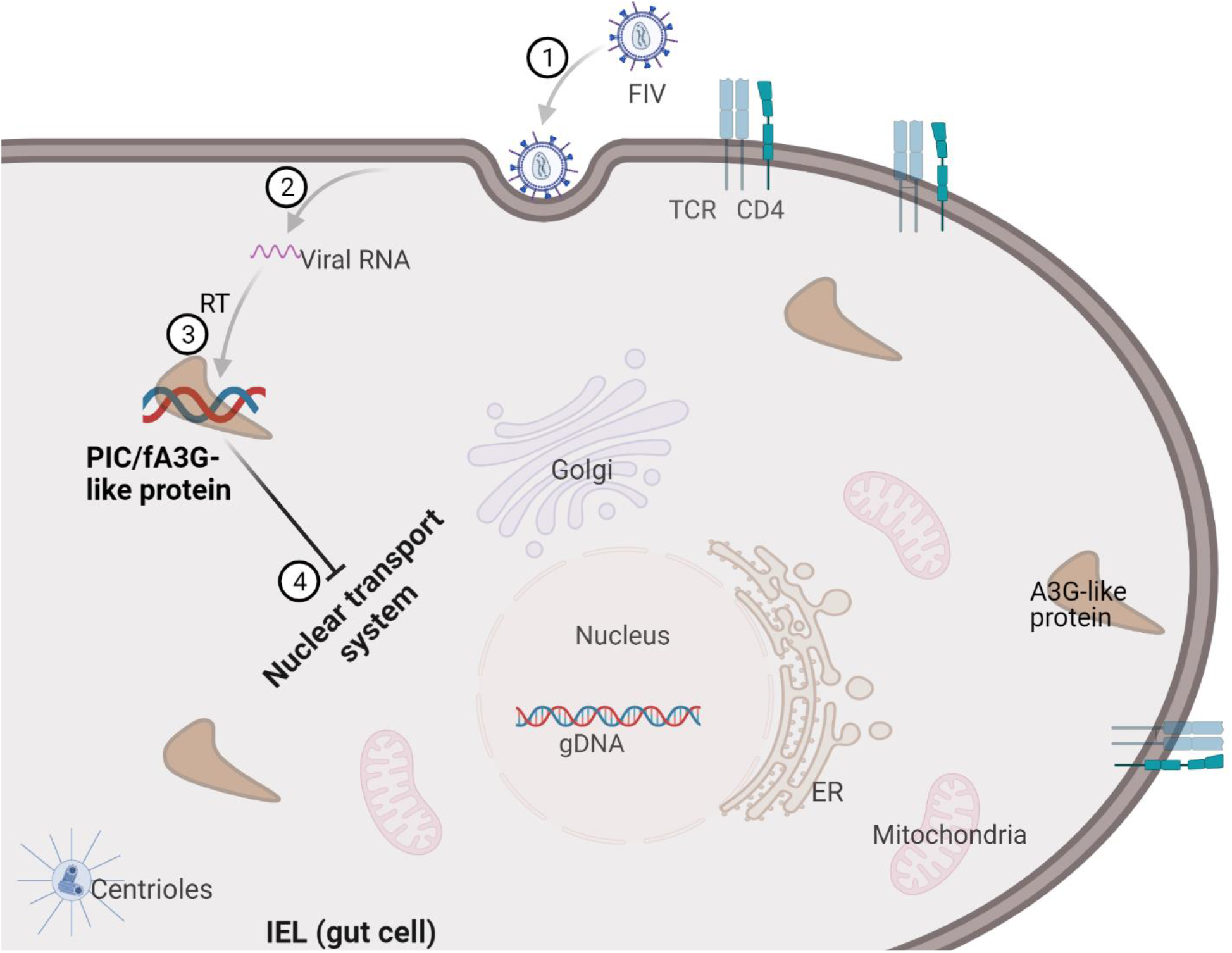
The proposed schematic diagram of the native fA3G-like protein determines the fate of the viral DNA in the cytoplasm. 1) at this stage, FIV breaks the integrity of the cell plasma membrane and delivers its genetic material into the cytoplasmic environment of the cell. 2) Viral RNA inside the hostile environment of the cell. 3) The viral RNA utilizes reverse transcriptase (enzyme) to achieve its reverse transcription to become viral DNA or pre-integration complex (PIC). At this stage, the native fA3G binds to the DNA and determines its fate by forcing PIC to remain inside the cytoplasm. The PIC has no other option than stay in the cytoplasm. This interaction could deteriorate once the cell gets activated.

## Acknowledgments

Plasmids (FIV-pF34, pF34/BH and pF34/3.4) were generous gifts from Dr. Ellen Sparger, UC Davis, School of Veterinary Medicine. Thanks to my students at KPU and UFV. Most experiments were conducted at The Ohio State University, and the principal author moved to Canada with the collected data. Partial financial support: National Institutes of Health and National Institute of Allergy and Infectious Diseases (Grants AI46220/KO2 AI50432).

## Author Contributions

Experiments design: Barnabe D. Assogba

Performed the experiments: Barnabe D. Assogba

Data analysis and organization: Barnabe D. Assogba, Raymond Soo, Jovy M.G. Assogba, Shannon Chaudhary, Harmanpreet Kaur, Mark A. P. Dela Cruz & Jovy M.G. Assogba.

Conference presentations: Shannon Chaudhary & Barnabe D. Assogba (The Annual Biomedical Research Conference for Minority Students & World Microbe Forum)

## References

1. Stremlau M, Owens CM, Perron MJ, Kiessling M, Autissier P, Sodroski J. The cytoplasmic body component TRIM5alpha restricts HIV-1 infection in Old World monkeys. Nature. 2004;427(6977):848–53.

2. Sheehy AM, Gaddis NC, Choi JD, Malim MH. Isolation of a human gene that inhibits HIV-1 infection and is suppressed by the viral Vif protein. Nature. 2002;418(6898):646–50.

3. Sayah DM, Sokolskaja E, Berthoux L, Luban J. Cyclophilin A retrotransposition into TRIM5 explains owl monkey resistance to HIV-1. Nature. 2004;430(6999):569–73.

4. Keckesova Z, Ylinen LMJ, Towers GJ, Coffin JM, Keckesova Z, Ylinen LMJ, et al. The Human and African Green Monkey TRIM5α Genes Encode Ref1 and Lv1 Retroviral Restriction Factor Activities. Proc Natl Acad Sci U S A. 2004;101(29):10780–5.

5. Hatziioannou T, Perez-Caballero D, Yang A, Cowan S, Bieniasz PD. Retrovirus resistance factors Ref1 and Lv1 are species-specific variants of TRIM5alpha. Proc Natl Acad Sci U S A. 2004;101(29):10774–9.

6. Yap MW, Nisole S, Lynch C, Stoye JP. Trim5alpha protein restricts both HIV-1 and murine leukemia virus. Proc Natl Acad Sci U S A. 2004;101(29):10786–91.

7. Holdeman R, Nehrt S, Strome S. A Single Amino Acid Change in the SPRY Domain of Human Trim5α Leads to HIV-1 Restriction. Current Biology. 2005;15(1):73–8.

8. Perez-Caballero D, Hatziioannou T, Zhang F, Cowan S, Bieniasz PD. Restriction of human immunodeficiency virus type 1 by TRIM-CypA occurs with rapid kinetics and independently of cytoplasmic bodies, ubiquitin, and proteasome activity. J Virol. 2005;79(24):15567–72.

9. Song B, Gold B, Colm O, Li X, Stremlau M, Winkler C, et al. The B30.2 (SPRY) Domain of the Retroviral Restriction Factor TRIM5 α Exhibits Lineage-Specific Length and Sequence Variation in Primates The B30.2 (SPRY) Domain of the Retroviral Restriction Factor TRIM5 ? Exhibits Lineage-Specific Length and Sequ. J Virol. 2005;79(10):6111–21.

10. Jarmuz A, Chester A, Bayliss J, Gisbourne J, Dunham I, Scott J, et al. An anthropoid-specific locus of orphan C to U RNA-editing enzymes on chromosome 22. Genomics. 2002;79(3):285–96.

11. Wedekind JE, Dance GSC, Sowden MP, Smith HC. Messenger RNA editing in mammals: New members of the APOBEC family seeking roles in the family business. Trends in Genetics. 2003;19(4):207–16.

12. Rogozin IB, Basu MK, Jordan IK, Pavlov YI, Koonin E V. APOBEC4, a New Member of the AID / APOBEC Family of Polynucleotide Report ND ES SC CE INTRODUCTION RIB. Cell Cycle. 2005;4(September):1281–5.

13. Konno Y, Nagaoka S, Kimura I, Yamamoto K, Kagawa Y, Kumata R, et al. New World feline APOBEC3 potently controls the inter-genus lentiviral transmission. Retrovirology. 2018;15(1).

14. Löchelt M, Romen F, Bastone P, Muckenfuss H, Kirchner N, Kim YB, et al. The antiretroviral activity of APOBEC3 is inhibited by the foamy virus accessory Bet protein. Proc Natl Acad Sci U S A. 2005;102(22):7982–7.

15. Russell R a, Wiegand HL, Moore MD, Schäfer A, Mcclure MO, Cullen BR, et al. Foamy Virus Bet Proteins Function as Novel Inhibitors of the APOBEC3 Family of Innate Antiretroviral Defense Factors Foamy Virus Bet Proteins Function as Novel Inhibitors of the APOBEC3 Family of Innate Antiretroviral Defense Factors. J Virol. 2005;79(14):8724–8731.

16. Powell LM, Wallis SC, Pease RJ, Edwards YH, Knott TJ, Scott J. A novel form of tissue-specific RNA processing produces apolipoprotein-B48 in intestine. Cell. 1987;50(6):831–40.

17. Honjo T, Nagaoka H, Shinkura R, Muramatsu M. AID to overcome the limitations of genomic information. Nat Immunol. 2005;6(7):655–61.

18. Conticello SG, Thomas CJF, Petersen-Mahrt SK, Neuberger MS. Evolution of the AID/APOBEC family of polynucleotide (deoxy)cytidine deaminases. Mol Biol Evol. 2005;22(2):367–77.

19. Mangeat B, Turelli P, Caron G, Friedli M, Perrin L, Trono D. Broad antiretroviral defence by human APOBEC3G through lethal editing of nascent reverse transcripts. Nature. 2003;424(6944):99–103.

20. Zheng Y hui, Irwin D, Kurosu T, Sata T, Peterlin BM, Tokunaga K. Human APOBEC3F Is Another Host Factor That Blocks Human Immunodeficiency Virus Type 1 Replication. J Virol. 2004;78(11):6073–6076.

21. Bishop KN, Holmes RK, Sheehy AM, Davidson NO, Cho SJ, Malim MH. Cytidine deamination of retroviral DNA by diverse APOBEC proteins. Current Biology. 2004;14(15):1392–6.

22. Lecossier D, Bouchonnet F, Clavel F, Hance AJ. Hypermutation of HIV-1 DNA in the Absence of the Vif Protein Author(s): Denise Lecossier, Francine Bouchonnet, François Clavel and Allan J. Hance Source: Science (1979). 2003;300(5622):1112.

23. Zhang H, Yang B, Pomerantz RJ, Zhang C, Arunachalam SC, Gao L. The cytidine deaminase CEM15 induces hypermutation in newly synthesized HIV-1 DNA. Nature. 2003;424(6944):94–8.

24. Wiegand HL, Doehle BP, Bogerd HP, Cullen BR. A second human antiretroviral factor, APOBEC3F, is suppressed by the HIV-1 and HIV-2 Vif proteins. EMBO J. 2004;23(12):2451–8.

25. Liddament MT, Brown WL, Schumacher AJ, Harris RS. APOBEC3F properties and hypermutation preferences indicate activity against HIV-1 in vivo. Current Biology. 2004;14(15):1385–91.

26. Stopak K, De Noronha C, Yonemoto W, Greene WC. HIV-1 Vif blocks the antiviral activity of APOBEC3G by impairing both its translation and intracellular stability. Mol Cell. 2003;12(3):591–601.

27. Sheehy AM, Gaddis NC, Malim MH. The antiretroviral enzyme APOBEC3G is degraded by the proteasome in response to HIV-1 Vif. Nat Med. 2003;9(11):1404–7.

28. Mehle A, Strack B, Ancuta P, Zhang C, McPike M, Gabuzda D. Vif Overcomes the Innate Antiviral Activity of APOBEC3G by Promoting Its Degradation in the Ubiquitin-Proteasome Pathway. Journal of Biological Chemistry. 2004;279(9):7792–8.

29. Chiu YL, Witkowska HE, Hall SC, Santiago M, Soros VB, Esnault C, et al. High-molecular-mass APOBEC3G complexes restrict Alu retrotransposition. Proc Natl Acad Sci U S A. 2006;103(42):15588–93.

30. Newman ENC, Holmes RK, Craig HM, Klein KC, Lingappa JR, Malim MH, et al. Antiviral function of APOBEC3G can be dissociated from cytidine deaminase activity. Current Biology. 2005;15(2):166–70.

31. Assogba BD, Leavell S, Porter K, Burkhard MJ. Mucosal administration of low-dose cell-associated feline immunodeficiency virus promotes viral latency. J Infect Dis. 2007;195(8):1184–8.

32. Dang Y, Wang X, Esselman WJ, Zheng YH. Identification of APOBEC3DE as another antiretroviral factor from the human APOBEC family. J Virol. 2006;80(21):10522–33.

33. Turelli P, Trono D. Editing at the crossroad of innate and adaptive immunity. Science. 2005;307(5712):1061–5.

34. Conticello SG, Harris RS, Neuberger MS. The Vif Protein of HIV Triggers Degradation of the Human Antiretroviral DNA Deaminase APOBEC3G. Current Biology. 2003;13(22):2009–13.

35. Kobayashi M, Takaori-Kondo A, Miyauchi Y, Iwai K, Uchiyama T. Ubiquitination of APOBEC3G by an HIV-1 Vif-Cullin5-Elongin B-Elongin C complex is essential for Vif function. Journal of Biological Chemistry. 2005;280(19):18573–8.

36. Oberste MS, Gonda MA. Conservation of amino-acid sequence motifs in lentivirus Vif proteins. Virus Genes. 1992;6(1):95–102.

37. Phillips TR, Talbott RL, Lamont C, Muir S, Lovelace K, Elder JH. Comparison of two host cell range variants of feline immunodeficiency virus. J Virol. 1990;64(10):4605–13.

38. Tomonaga K, Norimine J, Shin YS, Fukasawa M, Miyazawa T, Adachi A, et al. Identification of a feline immunodeficiency virus gene which is essential for cell-free virus infectivity. J Virol. 1992;66(10):6181–5.

39. Shacklett BL, Luciw PA. Analysis of the vif gene of feline immunodeficiency virus. Virology. 1994;204(2):860–7.

40. Jenkins Y, McEntee M, Weis K, Greene WC. Characterization of HIV-1 Vpr nuclear import: Analysis of signals and pathways. Journal of Cell Biology. 1998;143(4):875–85.

41. Vodicka M a., Koepp DM, Silver P a., Emerman M. HIV-1 Vpr interacts with the nuclear transport pathway to promote macrophage infection. Genes Dev. 1998;12(2):175–85.

42. Bukrinsky MI, Haggerty S, Dempsey MP, Sharova N, Adzhubei A, Spitz L, et al. A nuclear localization signal within HIV-1 matrix protein that governs infection of non-dividing cells. Nature. 1993;365(6447).

43. Gallay P, Hope T, Chin D, Trono D. HIV-1 infection of nondividing cells through the recognition of integrase by the importin/karyopherin pathway. Proc Natl Acad Sci U S A. 1997;94(18).

44. Popov S, Dubrovsky L, Lee MA, Pennathur S, Haffar O, Al-Abed Y, et al. Critical role of reverse transcriptase in the inhibitory mechanism of CNI-H0294 on HIV-1 nuclear translocation. Proc Natl Acad Sci U S A. 1996;93(21).

45. Dharan A, Talley S, Tripathi A, Mamede JI, Majetschak M, Hope TJ, et al. KIF5B and Nup358 Cooperatively Mediate the Nuclear Import of HIV-1 during Infection. PLoS Pathog. 2016;12(6).

46. Elhelu MA. The role of macrophages in immunology. J Natl Med Assoc y 1983 Mar;75(3):314-7 PMID: 6343621; PMCID: PMC2561478. 1983 Mar;75(3):314–7.

47. Mills CD, Kincaid K, Alt JM, Heilman MJ, Hill AM. M-1/M-2 Macrophages and the Th1/Th2 Paradigm. The Journal of Immunology. 2000;164(12).

48. Chiu YL, Witkowska HE, Hall SC, Santiago M, Soros VB, Esnault C, et al. High-molecular-mass APOBEC3G complexes restrict Alu retrotransposition. Proc Natl Acad Sci U S A. 2006;103(42).

49. Assogba BD, Leavell S, Porter K, Burkhard MJ. Mucosal administration of low-dose cell-associated feline immunodeficiency virus promotes viral latency. Journal of Infectious Diseases. 2007;195(8).

50. Shukla E, Chauhan R. Host-HIV-1 Interactome: A Quest for Novel Therapeutic Intervention. Cells. 2019;8(10).

